# Epigenenomic and transcriptomic characterization of *NPM1*-mutated CN-AML subtypes

**DOI:** 10.1101/2024.05.07.592738

**Authors:** Lia N’Guyen, Clara Tellez-Quijorna, Julien Vernerey, Sylvain Garciaz, Nadine Carbuccia, Arnaud Guille, Claire Burny, Mathilde Poplineau, Anne Murati, Marie-Anne Hospital, Norbert Vey, Daniel Birnbaum, Estelle Duprez

## Abstract

Cytogenetic normal AML with *NPM1* mutations forms a distinct AML entity, associated with an intermediate prognosis and a heterogeneous response to treatment. We previously described an epigenetic biomarker, defined by the level of H3K27me3 on 70kb of the *HIST1* cluster in patient blast DNA. This epigenetic mark separates cytogenetically normal *NPM1*^mut^ AML into two groups of patients differing in their survival rate following chemotherapy. To better characterize the influence of the biomarker on disease progression, we performed transcriptomic and histone mark profiling on patient blasts according to the level of H3K27me3 *HIST1*. Our integrated analysis revealed that the two groups of patients display differences in terms of transcriptomic, chromatin landscape and cell surface markers, which could explain the clinical difference. Our profiling revealed novel targets and therefore constitutes an (epi)transcriptomic resource for *NPM1* AML. It also highlights the power of epigenetic profiling to dissect the heterogeneity of a single AML genetic entity.

## INTRODUCTION

Acute Myeloid leukemia (AML) is a severe hematological disorder characterized by molecular and cellular heterogeneity. The various cytogenetic and molecular abnormalities identified in AML have major repercussions in terms of prognosis and treatment and have fueled precision medicine ^1^. One of the most frequent genetic lesions found in AML is the mutation in the *NPM1* gene, being detected in around 50-60% of AMLs with a normal karyotype (CN-AMLs) ^2^ and for which prevalence and prognostic impact have been extensively reported ^3,4^. In the revised 2016 World Health Organization classification *NPM1*^mut^ AML was recognized as a distinct entity associated with an intermediate prognosis and heterogeneity of outcome and response to treatment ^5,6^.

The unique feature of *NPM1*^mut^ AML relies on NPM1 functions. NPM1 protein is a multifunctional protein mainly involved in the regulation of ribosomal protein assembly and trafficking, DNA repair process, genome stability, and gene regulation by altering chromatin structure ^2^. All of these functions can be impaired by the abnormal localization of the NPM1^mut^ protein, resulting in a nuclear-to-cytoplasmic translocation that contributes to leukemogenesis ^7^. Recently, the mutant form of NPM1 has been shown to have a neomorphic function by hijacking and amplifying the AML transcriptional program at H3K27ac-enriched genes ^8,9^. This epigenetic gain-of-function is consistent with the transcriptional profiles associated with *NPM1*^mut^ AML ^10^ and, above all, supports the development of epigenetic therapies aimed at targeting AML cells with mutant NPM1^11^.

The heterogeneity observed within this entity can be explained by the co-occurrence of additional mutations, such as *FLT3*^ITD^ and *DNMT3A*^mut^, which influence prognosis and form the basis of genetic-based risk stratification models ^12,13^. Although co-mutational patterns with prognostic implications are regularly updated with machine learning approaches ^1^, they remain insufficient to avoid diagnostic and therapeutic pitfalls in *NPM1*^mut^ AML ^14^. The search for and identification of new types of biomarkers, such as epigenetic signatures, which reflect cellular identity and dynamics are relevant in *NPM1*^mut^ AML. Following this idea, we previously identified an epigenetic biomarker, defined as a H3K27me3 enrichment on 70 Kb of the *HIST1*cluster on chromosome 6, associated with *NPM1* mutation ^15,16^. This epigenetic biomarker impacted patient outcome and discriminated two groups of patients (H3K27me3*HIST1*^low^ and H3K27me3*HIST1*^high^ herein named Me3*HIST1*^low^ and Me3*HIST1*^high^) in the cytogenetically normal *NPM1*^mut^ (CN-*NPM1*^mut^) AML entity.

To progress in the understanding of the function and mechanism of *NPM1*^mut^ in link with epigenetic regulation, we combined high-quality ChIP-seq and RNA-seq across CN-*NPM1*^mut^ AMLs to comprehensively characterize the epigenetic, transcriptomic and cellular consequences associated with and determining the classification of the two Me3*HIST1*^low^ and Me3*HIST1*^high^ subtypes of this entity.

## MATERIAL and METHODS

### Patient samples

A total of 35 CN-*NPM1*^mut^ AML samples, with at least 62% of blasts were selected from AML samples stored at Institut Paoli-Calmettes (IPC) Tumor Bank at diagnosis. In this tumor bank, blast cells are separated from blood or bone marrow (BM) samples through density-gradient (ficoll) separation and stored in liquid nitrogen. Informed consent was provided by all patients according to the Declaration of Helsinki and subjected to ethical institutional review board approval. All samples have undergone mutational analysis (custom sequencing-based assay) to assess for frequent *NPM1*^mut^ associated mutations (*IDH1,2, FLT3*ITD and *DNMT3A*).

Cell surface marker expression was performed at diagnosis by the cytology platform of the IPC. Most of samples were subjected to ChIP-seq (n=24; four histone marks) and RNA-seq (n=19).

### Chromatin immunoprecipitation (ChIP-seq)

For histone mark ChIP-seq, cells from BM or blood aspirate were fixed with 1% of formaldehyde for 8 min. Reaction was quenched by adding 2 mM of glycine. ChIP were performed as previously described procedures ^17^, using 7.5 µg of chromatin per histone marks. Briefly, the chromatin of interest was immunoprecipitated together with 0.75 µg spike-in drosophilia chromatin overnight at 4°C with magnetic beads pre-incubated with the following antibodies: H3K27me3 (Cell Signalling #9733S), H3K27ac (Active Motif #39133), H3K4me1 (Abcam #ab8895), H3K4me3 (Abcam #ab8580). Immunoprecipitates were washed and immunoprecipitated DNA was purified with I-Pure Kit V2 (Diagenode). ChIP-seq libraries were generated using the MicroPlex Library Preparation Kit (Diagenode) following the manufacturer’s instructions and analyzed on a 2100 Bioanalyzer system (Agilent) prior sequencing. Sequencing was performed with a NextSeq500 sequencer (Illumina) using a 75-nt single-end protocol, at the Paoli Calmettes Institute Sequencing Facility (IPC, Marseille).

#### Alignment

Sequencing quality control was determined using the FastQC tool (http://www.bioinformatics.babraham.ac.uk/projects/fastqc/) and aggregated across samples using MultiQC. Reads with a Phred quality score less than 30 were filtered out. Reads were mapped to the human (Hg38) reference genome using default parameters of Bowtie2 (v2.3.4.3) ^18^. Only mapped reads were filtered in using samtools (v1.9) (*view; --exclude-flags 4*). Duplicate tags were removed using Picard Tools (v2.20.2) (https://broadinstitute.github.io/picard/) (*MarkDuplicates*; *REMOVE_DUPLICATES = true*) and mapped tags were processed for further analysis.

#### BigWigs

Mapped reads were normalized by their library size and converted into bigWigs using the deepTools suite (v3.2.1) ^19^ (*bamCoverage --scaleFactor library_size_factor --binSize 5*). Reads were extended using values previously computed by deepTools (*bamPEFragmentSize*) and used through the *--extendReads* option. BigWig files from multiple replicates were merged into a single file using a custom bash script (*bigWigMerge*). All resulting files were visualized using the Integrative Genomics Viewer (IGV; v2.5.2)

#### Peak calling

High confidence binding sites were determined using MACS2 callpeak (v2.1.2) ^20^ in broad mode *(--broad broad-cutoff=0*.*05 -q 0*.*01 --nomodel*). Each sample’s fragment size was previously computed by deepTools (*bamPEFragmentSize*) and used through the *--extsize* option. Inputs were used as controls.

#### Differential analysis

Initially, a consensus peak set was determined among samples using MACS2 output beds with bedtools (v2.28.0) ^21^ (*genomecov -bg -g chrominfo)*. Coordinates shared by at least 2 samples (i.e. coverage >= 2) were selectively retained and merged into unified coordinates (*bedtools merge*). Mapped tags were then counted within these consensus coordinates using featureCounts (v1.6.4) ^22^ and normalized using DESeq2 (*estimateSizeFactors*) (v1.26.0) ^23^. Finally, differential analysis was performed (*DESeq, minReplicatesForReplace = Inf, betaPrior = T*).

#### Enhancers

BEDtools intersect (v2.17.0) with a minimum overlap of 1-base was used to determine enhancer coordinates. Overlap significances were computed by hypergeometric tests (phyper function, lower.tail=False). To map H3K27ac and H3K27me3 signals on enhancer coordinates we applied computeMatrix (v3.5.0) from the deepTools suite (*--afterRegionStartLength -- beforeRegionStartLength --referencePoint = center –skipZeros)*.

All H3K27ac peaks within ± 2.0 kb around transcription start sites (TSSs) were first excluded. The remaining peaks closer than default distance of 12.5 kb were stitched together, and subsequently ranked by normalized H3K27ac level corrected by input background. Finally, Enhancers were separated from typical enhancers based on the inflection point of H3K27ac signal curve. We defined 3 groups of enhancers according to their epigenetic marks: active (H3K4me1 + H3K27ac), poised (H3K4me1 + H3K27me3) or intermediate (H3K4me1). Differential enhancer activities between AML subtypes were identified using DESeq2 ^23^ with an adjusted pValue less than 0.1 and absolute fold change greater than 1.5. Enhancer assignment to the nearest genes was determined by BEDTools ^21^.

#### H3K4me3 signal analysis

H3K4me3 signals from AML patients were analyzed using a custom R (version?) script. Briefly, H3K4me3ChIP-Seq differential analysis were annotated using both Homer (v4.11) ^24^ (*annotatePeaks*.*pl*) and ChIPseeker (v1.34.0) ^25^ (*annotatePeak*) annotation tools. Only significant differential peaks were kept (pValue < 0.05) and split into 3 groups:

- TSS group, including all peaks located within 1kb around the TSS.
- Intragenic group, including peaks annotated as 5UTR, Exon, Intron, 3UTR, TTS, or non-coding.
- Intergenic group, included all other peaks.

Only TSS and intragenic peaks were retained for further analysis. H3K4me3 TSS signals (log2FC) and H3K4me3 Intragenic signals (log2FC) were plot against each other using ggplot2 (v3.4.1). Peaks were highlighted into 4 sub-grouped (i.e. TSSup INTRAup, TSSdown INTRAdown, TSSup INTRAdown, TSSdown INTRAup) regarding their respective directionality between Me3*HIST1*^high^ and Me3*HIST1*^low^ patients.

### RNA-sequencing

Total RNA was isolated from 5 × 10^6^ cells harvested from frozen patient vials using the RNeasy Plus mini-Kit (Qiagen). Libraries were prepared with the NEBNext® rRNA Depletion Kit and were sequenced using the Illumina NextSeq500 platform, 75-bp paired-end reads, at 50 million reads per sample at the HELIXIO platform.

#### Alignment

Sequencing quality control was determined using the FastQC tool (http://www.bioinformatics.babraham.ac.uk/projects/fastqc/) and aggregated across samples using MultiQC (v1.7) ^26^. Reads with a Phred quality score less than 30 were filtered out and mapped to the human (Hg38) reference genome using Subread-align (v1.6.4) with default parameters and Gene expression quantification was performed by counting mapped tags at gene levels using featureCounts (v1.6.4) ^22^.

#### Differential analysis

Differentially expressed genes between the 8 Me3*HIST1*^low^ *versus* the 11 Me3*HIST1*^high^ samples were identified using the Bioconductor package DESeq2 (v1.26.0) ^23^ (*DESeq, minReplicatesForReplace = Inf, betaPrior = T*) and were selected according to the following criteria: pValue<0.05, absolute log2FC >1.

#### Functional enrichment analysis

The Gene Ontology (GO) analysis was performed using clusterProfiler with a pValue<= 0.05 and by considering a background. Then, the genes were split in two groups according to their fold-change (UP genes: log2(FC) >= 1, DOWN genes: log2(FC) <= 1). The GO categories chosen were Biological Process (BP) and KEGG pathways. Volcano plots, GO enrichment and Cnetplots figures were plotted by https://www.bioinformatics.com.cn/en, a free online platform for data analysis and visualization. For the gene set enrichment analysis (GSEA), we established a list of 3794 genes using a nominal p-value threshold of 0.05 corresponding to an adjusted p-value threshold of 0.28. Genes were then ranked to both their statistical significance by using the Log10(p-value) × sign of the Log2(FC) as input of the GSEA software (v4.2.2). Specific gene sets for the GMP, CSH and MEP populations were generated from HSPC data from healthy donors ^27^ (GSE183252). For each of these 3 populations, only genes displaying an enriched expression of 2-fold compared to the other 2 populations were retained.

### Multiomic analysis

Output peaks from H3K27ac ChIP-Seq differential analysis were annotated using both Homer (v4.11) ^24^ (*annotatePeaks*.*pl*) and ChIPseeker (v1.34.0) ^25^ (*annotatePeak*) annotation tools. Bulk RNA-seq and ChIP-seq data were then merged based on their gene names.

Only candidates with a significant differential in both ChIP-seq and RNA-seq (pValue < 0.05 and |Log2FC| *≥* 1, represented by dotted lines) were plotted against each other using ggplot2 (v3.4.1). The top 10 candidates per quadrant were labelled using ggrepel (v0.9.3) (*geom_text_repel*).

### Statistics

The statistical significance of the overlap among any peak or gene sets was determined by the hypergeometric test. Comparison between different groups was tested using Fisher’s exact test for dichotomous variables and Mann-Whitney test for continuous variables. Differential expression analysis were done with DESeq2.

## DATA AVAILABILITY

The CHIP-seq and RNA-seq data generated here are available in the Gene Expression Omnibus database under accession code GSE262473.

H3K27me3 CHIP-seq BigWigs to generate figure 4E were downloaded from the https://epigenomesportal.ca/ihec/grid.html

## RESULTS

### Me3*HIST1* ^low^ and Me3*HIST1*^high^ NPM1 AMLs show differences

To characterize Me3*HIST1*^low^ and Me3*HIST1*^high^ *NPM1*^mut^ patients, we selected CN-*NPM1*^mut^ 35 AML samples from our biobank and chose to investigate the chromatin landscape and the RNA expression in 24 (9 Me3*HIST1*^low^ and 15 Me3*HIST1*^high^) and 19 (8 Me3*HIST1*^low^ and 11 Me3*HIST1*^high^) samples respectively (**Figure 1A**).

**Figure 1:**
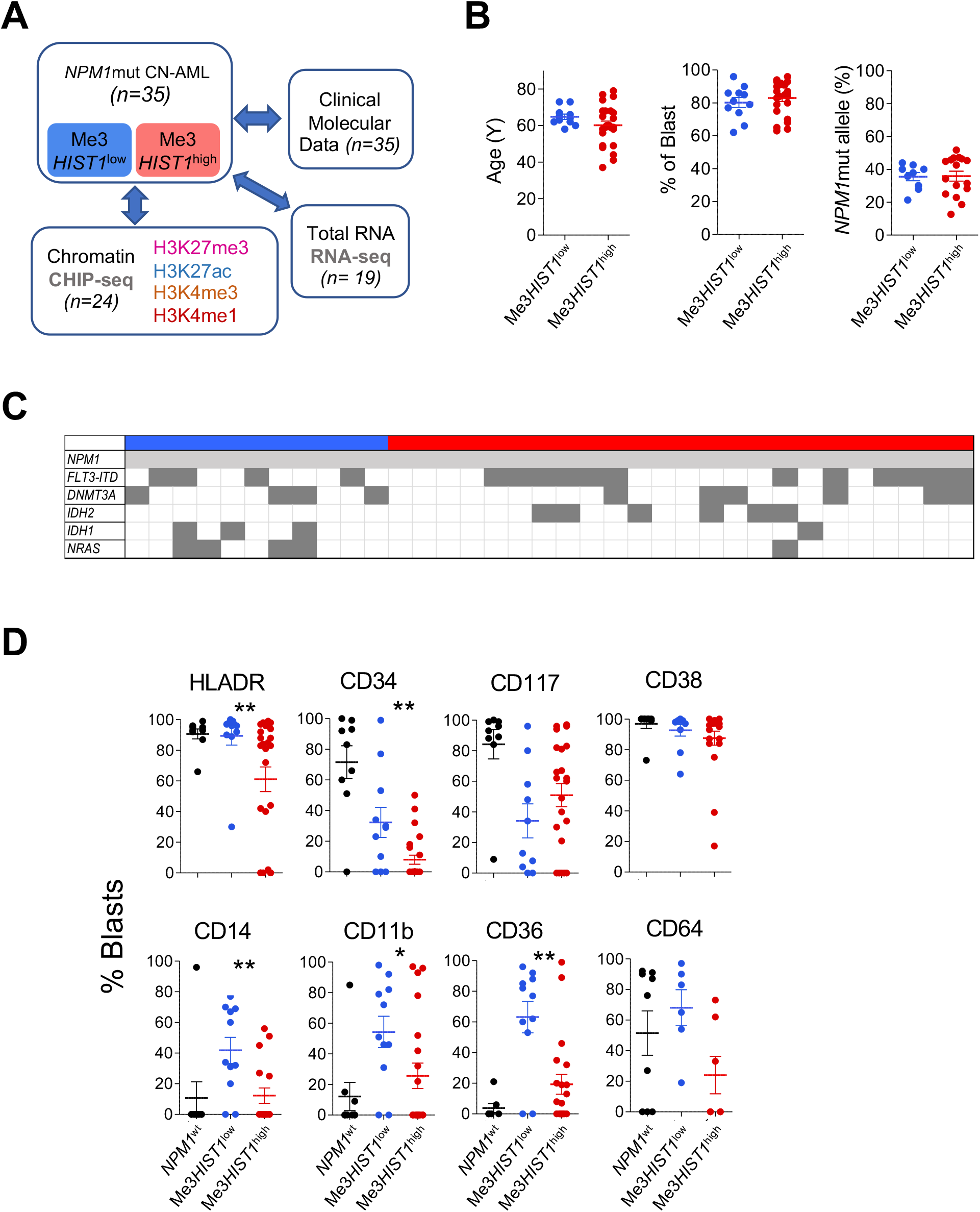
Characterization of 2 subgroups of *NPM1*-mutated CN-AML. (**A**) Experimental scheme and patient presentation. (**B**) Age, percentage of blasts and *NPM1* mutated allele frequency distribution in the two AML subgroups. (**C**) Occurrence of mutations frequently associated with *NPM1*^mut^ according to Me3*HIST1*^high^ and Me3*HIST1*^low^. (**D**) Percentage of HLADR, CD34, CD117, CD38, CD14, CD11b, CD36 and CD64 positive blasts according to *NPM1* mutation and Me3*HIST1* status. (** p<0.01)

As previously reported, we confirmed the absence of difference in age at diagnosis or blast percentage between the two groups (**Table 1, Figure 1B, left-middle panels**). We also verified that the *NPM1* mutated allele frequency, which influences patient outcome ^28^, was similar in our two groups of patients (**Table 1, Figure 1B, right panel**). Distribution of the most frequent co-occurring mutations *DNMT3A*^mut^ or *FLT3*^ITD^ was equal between the two groups, while *IDH2*^R140^ had the tendency to be overrepresented in *NPM1*^mut^ Me3*HIST1*^high^ subgroup (25% versus 0%, *p=0*.*15*) as previously observed ^16^. We also observed in this cohort an overrepresentation of *IDH1* and *RAS* mutations in CN-*NPM1*^mut^ Me3*HIST1*^low^ (27.27% vs 4.17%, *p=0*.*082* and 36.36 vs 4.17%, *p=0*.*026* respectively) (**Table 1, Figure 1C**). However, this overrepresentation was not confirmed when analyzing a bigger cohort (N=100) (data not shown). The two groups showed some immunophenotypic differences. Me3*HIST1*^low^ patients presented significantly higher levels of CD34 and HLADR compared to Me3*HIST1*^high^ patients, while they presented no difference in the expression level of immature markers CD38 or CD177 (**Table 1, Figure 1D**). Interestingly, we also observed a higher percentage of CD14-, CD11b- and CD36-positive blasts in the Me3*HIST1*^low^ group without detecting any difference in CD64 (**Table 1, Figure 1D**). These immunophenotypic profiles are indicative of a more monocytic leukemia phenotype that is associated with a specific energy metabolism in the Me3*HIST1*^low^ group.^29, 30^

**Table 1:** patient characteristics.

| Characteristics |  | All patients<br>(n=35) | Me3 HIST1<sup>low</sup><br>(n=11) | Me3 HIST1<sup>high</sup><br>(n=24) | P |
| --- | --- | --- | --- | --- | --- |
| Age, y |  |  |  |  |  |
|  | Median | 62 | 64 | 59.5 | 0.20 |
|  | Range | 37-77 | 58-73 | 37-79 |  |
| Sex, % |  |  |  |  |  |
|  | Male |  | 9.1 | 46 |  |
| Blast, % |  |  |  |  |  |
|  | Median | 85 | 81 | 85.5 | 0.46 |
|  | Range | 62-96 | 62-96 | 63-96 |  |
| Molecular alterations, % |  |  |  |  |  |
| VAF-NPM1mutant |  | 38.2 | 37.9 | 38.5 |  |
|  | FLT3ITD | 45.71 | 36.36 | 50 | 0.49 |
|  | DNMT3A | 28.57 | 36.36 | 25 | 0.69 |
|  | IDH2 (R140) | 17.14 | 0 | 25 | 0.15 |
|  | IDH1 (R132) | 11.43 | 27.27 | 4.17 | 0.082 |
|  | N-RAS | 14.29 | 36.36 | 4.17 | 0.026 |
| Expression, % |  |  |  |  |  |
| HLADR | Mean |  | 89.45 | 61.08 | 0.0080 |
|  | Range | 0-100 | 30-100 | 0-99 |  |
| CD34 | Mean |  | 32.27 | 7.96 | 0.0088 |
|  | Range | 0-99 | 0-99 | 0-50 |  |
| CD117 | Mean |  | 34.1 | 50.91 | 0.253 |
|  | Range | 0-97 | 0-96 | 0-97 |  |
| CD38 | Mean |  | 92.60 | 87.52 | 0.3062 |
|  | Range | 17-100 | 64-100 | 17-100 |  |
| CD14 | Mean |  | 41.82 | 12.29 | 0.0020 |
|  | Range | 0-91 | 0-77 | 0-91 |  |
| CD11b | Mean |  | 54.27 | 25.60 | 0.0439 |
|  | Range | 0-98 | 0-98 | 0-97 |  |
| CD36 | Mean |  | 63.18 | 19.43 | 0.0075 |
|  | Range | 0-99 | 0-96 | 0-99 |  |
| CD64 | Mean |  | 68 | 51.44 | 0.636 |
|  | Range | 0-97 | 19-97 | 0-92 |  |

### Epigenetic differences separated *NPM1*^mut^ CN-AMLs

Because chromatin and epigenetic dynamics have the potential to impact AML progression ^31^, we profiled four histone marks H3K27ac, H3K27me3, H3K4me1 and H3K4me3 according to Me3*HIST1*^low^ and Me3*HIST1*^high^ status by ChIP-sequencing. Zoom on the *HIST1* locus confirmed the difference in H3K27me3 level on a part of the *HIST1* cluster covering 70 kb and 11 histone genes (**Figure S1**) that distinguished Me3*HIST1*^low^ and ^high^ patients. IGV visualization (**Figure S2**) and bin quantification (**Figure 2A**) confirmed the localized and broad nature of the H3K27me3 chromatin domain that defined the Me3*HIST1*^high^ AML. Mirroring this H3K27me3 chromatin domain, the H3K27ac signal was weakly detected in the *HIST1* region of the Me3*HIST1*^high^ group compared to the Me3*HIST1*^low^ group (**Figure 2A; Figure S2A**). In addition, a comparison of the H3K9me3 mark, which is involved in heterochromatin repression across 4 Me3*HIST1*^low^ and 4 Me3*HIST1*^high^ cases confirmed the repressive state of the *HIST1* region delineated with H3K27me3 enrichment in Me3*HIST1*^high^ patients (**Figure 2B**). H3K27ac and H3K4me3 signals in Me3*HIST1*^low^ patients were characterized by fairly sharp peaks positioned in intragenic regions (**Figure 2C; Figure S2A)**. The specificity of this epigenetic profile at the *HIST1* locus was confirmed by analysis of another histone locus, *HIST2* (**Figure S2B&C**), for which H3K27me3 enrichment was never detected. Thus, we confirmed and underlined the specificity of Me3*HIST1*^high^ and Me3*HIST1*^low^ status by ChIP-seq and highlighted a link with repressive marks and Me3*HIST1*^high^ at the *HIST1* locus in CN-*NPM1*^mut^ AMLs.

**Figure 2:**
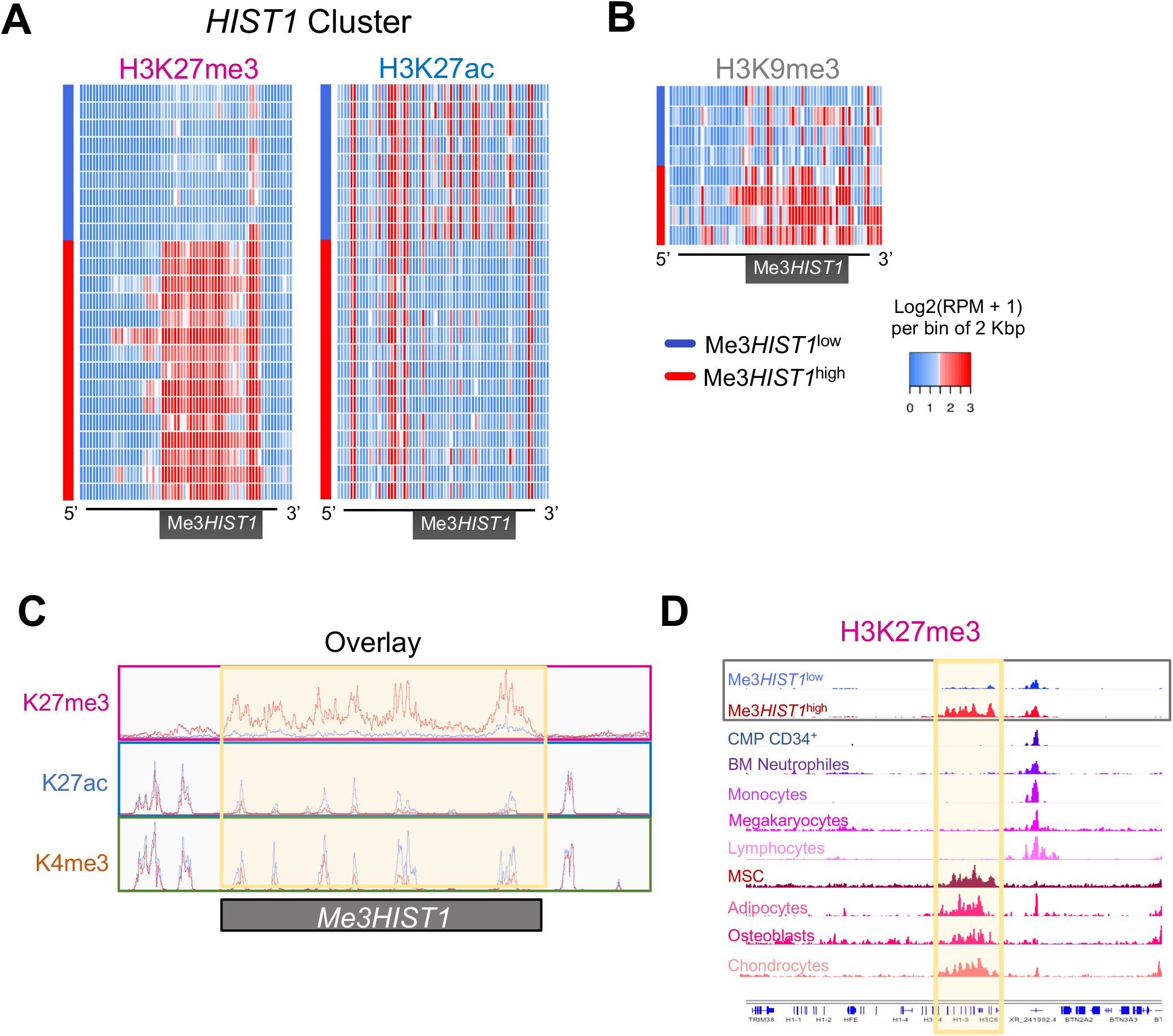
H3K27me3*HIST1* enrichment is anti-correlated with active marks. (**A-B)** Quantification and representation of normalized ChIP-seq signals on part of *HIST1* Cluster (Chr6 26160000-26300000). H3K27me3*HIST*^high^and^low^groups of patients are indicated by stacked red and blue bars respectively. One lane represents one patient. ChIP-seq signal is averaged by bins of 2kbp and 1 bin expressed in Log2(RPM + 1) unit. In **(A)** H3K27me3 and H3K27ac signal are vizualized in 24 patients and in (**B**), H3K9me3 signal is represented in 8 patients. (**C)** Overlay of the H3K27me3, H3K27ac and H3K4me3 signal averaged from the 9 Me3*HIST1*^low^ (blue line) and 15 Me3*HIST1*^high^ (red line) patients. (**D**) Visualization of Me3*HIST1*^high^ profiles though hematopoietic and mesenchymal lineages using public data. H3K27me3*HIST1* cluster region is highlighted in yellow.

To assess whether Me3*HIST1*^high^ could be found in physiological conditions, we screened H3K27me3 signal through the *HIST1* locus across hematopoietic differentiation using publicly available datasets. We did not observe enrichment in any of the hematopoietic lines tested, suggesting a gain of H3K27me3 irrespective of leukemia differentiation stage (**Figure 2D**). Interestingly, we detected the same H3K27me3 domain in bone marrow-derived mesenchymal stem cells (MSCs) as well as in their progeny (**Figure 2D)**, suggesting that the H3K27me3*HIST1* domain is present under physiological differentiation lineages.

No difference was observed in the global signal of any of H3K27ac, H3K27me3, H3K4me1 and H3K4me3 marks between the two groups of patients (**Figure S3A)**. However, differential analysis of the genomic distribution of the four histone marks between the two patient groups revealed that the H3K4me3 signal was less abundant at the transcription start site (TSS) in Me3*HIST1*^high^ patients, but increased at intragenic regions compared to Me3*HIST1*^low^ patients (**Figure 3A**). This H3K4me3 redistribution did not correspond to a sharp-to-broad peak profile change but to a decrease in TSS H3K4me3 peak signal (**Figure S3B)**. We identified 52 genes that significantly lost TSS H3K4me3 signal with 20 of them gaining intragenic H3K4me3 signal in Me3*HIST1*^high^ patients in majority related to metabolism (**Figure 3B&C; Table S1)**. Signal visualization confirmed the decrease of H3K4me3 in TSS counterbalanced by an increase in intragenic signal (**Figure 3D**), which could influence PolII progression ^32^.

**Figure 3:**
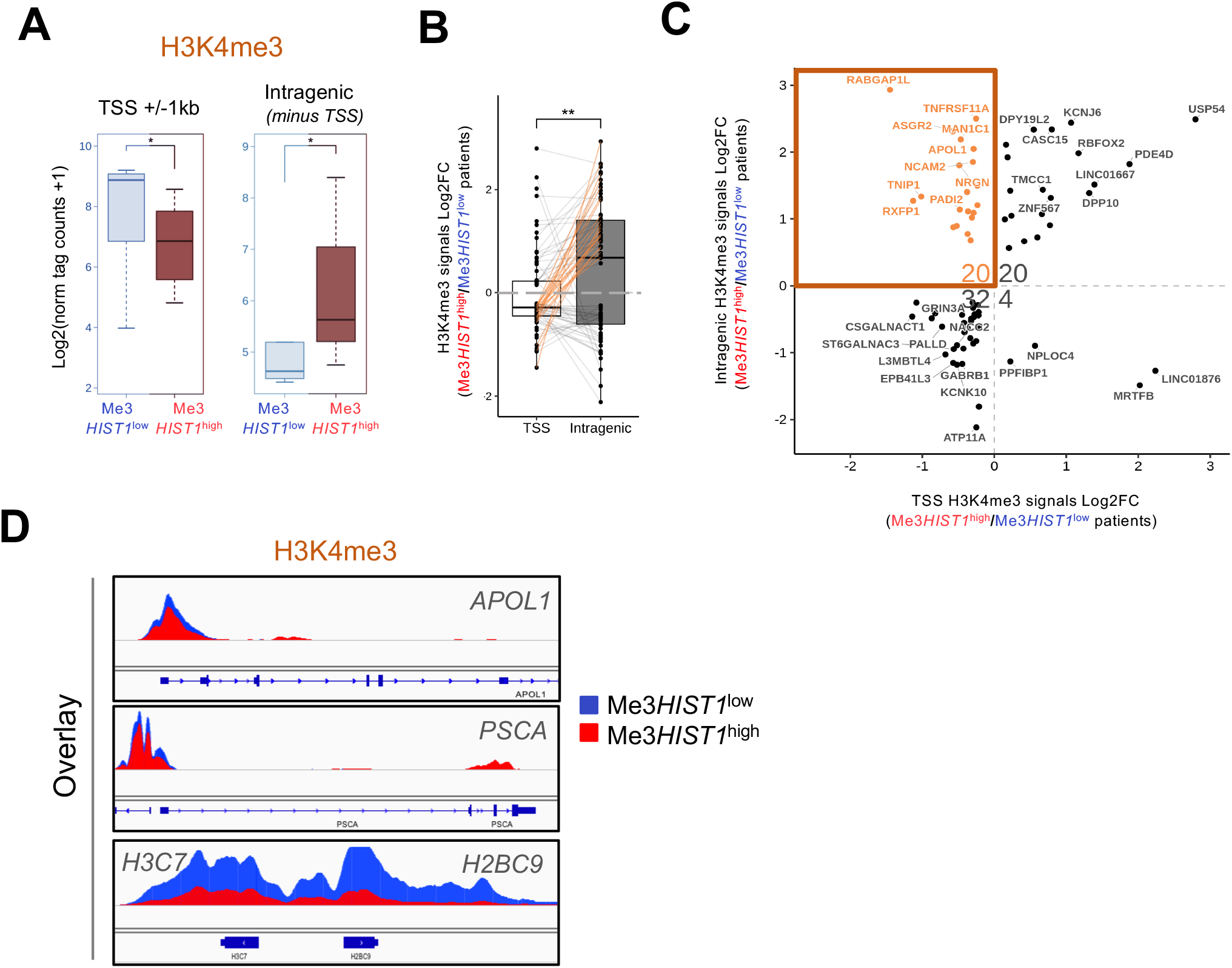
H3K27me3*HIST1* enrichment has subtle influence on histone mark level and distribution. (**A)** Comparison of the genome-wide H3K4me3 signal between the 9 Me3*HIST1*^low^ (blue line) and 15 Me3*HIST1*^high^(red line) at TSS and intragenic regions. Signal is express as a normalized tag counts (*p<0.05). **(B)** Boxplots of the differences in H3K4me3 signal between Me3*HIST1*^low^and Me3*HIST1*^high^ patients across TSS and intragenic regions (** p<0.01). A line connects both TSS and intragenic measured peaks of a gene. The orange lines highlight a sub-population of genes (see in **C**). (**C)** Identification of genes presenting significant H3K4me3 signal differences at TSS and intragenic regions between Me3*HIST1* high and Me3*HIST1*^low^ patients. The scatterplot represents the Log2FC of the Me3*HIST1* high /Me3*HIST1*^low^ ratio of H3K4me3 signal at TSS (x axis) and intragenic (y axis). (**D)** Overlay of the averaged H3K4me3^low^ signal obtained from the 9 Me3*HIST1* (blue) and 15 Me3*HIST1*high (red) patients zoomed at *APOL1, PSCA* and *H3C7-H2BC9* genomic localizations.

### Me3*HIST1*^low^ and Me3*HIST1*^high^ shape transcriptomic signatures in *NPM1*mut CN-AML

Differentially expressed gene (DEG) analysis of RNA-seq data from the patient blasts identified 372 genes significantly UP in Me3*HIST1*^low^ condition and 656 genes UP in Me3*HIST1*^high^ (**Figure 4A; Table S2)**. The Me3*HIST1*^low^ upregulated gene set was enriched in chromatin and nucleosome assembly while the Me3*HIST1*^high^ upregulated gene set was enriched in hemostasis and primitive hematopoiesis (**Figure 4B)**. In Me3*HIST1*^low^, the chromatin functional enrichment was driven by the upregulation of the histone genes covered by the signature (**Figure S4A)**. To better characterize the two groups, we interrogated different signatures through gene set enrichment analysis (GSEA). We found that Me3*HIST1*^low^ AML presented a strong correlation with E2F signature and cell proliferation, both hallmarks of unfavorable progression and in line with our previous data ^16^. We also revealed that the Me3*HIST1*^high^ AML harbored a high ribosomal protein signature and was correlated with previous published CN-*NPM1*^mut^ signatures^10, 33^ (**Figure 4C; Figure S4B)**. Some genes signing *NPM1*^mut^ such as *FOXC1, HOXB* were also markers of the different biological processes that characterized Me3*HIST1*^high^ (**Figure S4C)**. We also tested different signatures reflecting hematopoietic cell maturation. Consistent with the immunophenotype, we found that Me3*HIST1*^low^ leukemia was more committed to the myeloid lineage with enrichment for monocyte, neutrophil and GMP genes, whereas Me3*HIST1*^high^ leukemia presented a more pronounced megakaryocyte potential with enrichment for MkP, MEP and HSC signatures (**Figure 4D; Figure S4D)**. We also found that Me3*HIST1*^low^ blasts were enriched with an established CD36 signature ^30^ (**Figure 4E; Table S3**). By intersecting up-regulated genes in Me3*HIST1*^low^ with CD36 signature genes and down-regulated genes in NPM1^mut^ AMLs, we unveiled that the *KYNU* gene encoding the Kureinin enzyme involved in tryptophan metabolism and immunomodulation ^34^ was up-regulated in Me3*HIST1*^low^ AMLs.

**Figure 4:**
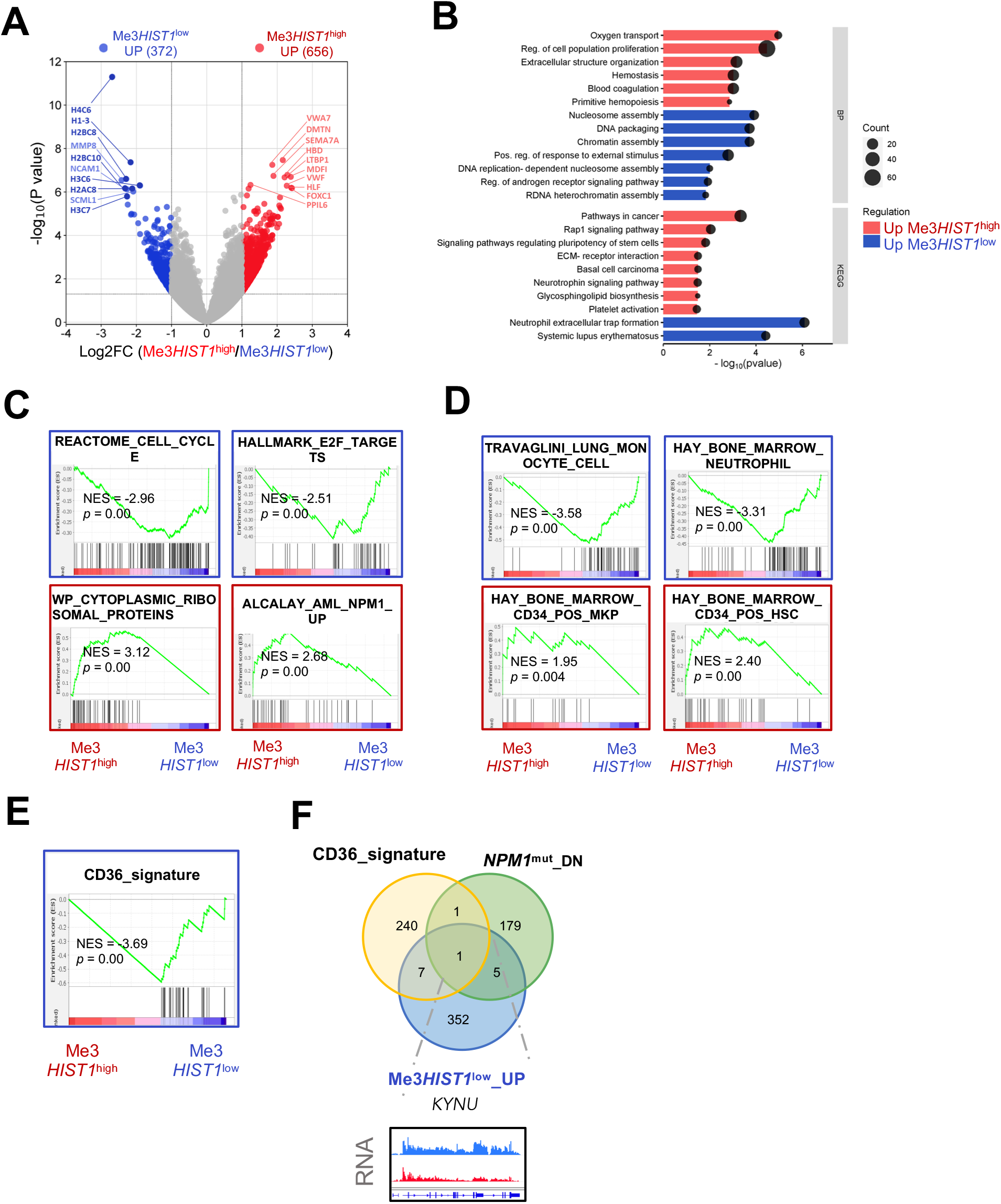
Me3*HIST1*^high^ AML is more megakaryocytic and less proliferative than Me3*HIST1*^low^ AML. (**A)** Volcano plot of gene expression changes between Me3*HIST1*^low^ and Me3*HIST1*^high^. The 10 most significantly upregulated genes are labelled in blue (Me3*HIST1*^low^ UP) or in red (Me3*HIST1*^high^ UP), and candidates are color-coded using thresholds of |Log2FC|=1 and nominal p-value < 0.05. **(B)** Functional enrichment analysis performed separately for the up-regulated genes in Me3*HIST1*^low^ and Me3*HIST1*^high^ conditions from GO (gene ontology) BP (biological process) and KEGG databases. The dot size represents the number of genes. The blue and red color indicates the Me3*HIST1*^low^ Up and Me3*HIST1*^high^ Up directionality, respectively. (**C-D)** Gene set enrichment analysis of Me3*HIST1*^low^ and Me3*HIST1*^high^ AMLs against: in (**C**) cell cycle (REACTOME), E2F targets (HALMARK), cytoplasmic ribosomal proteins and NPM1^mut^ AML (ALCALAY) and in (**D**) hematopoietic cell gene sets (MONOCYTE, NEUTROPHIL, MKP and HSC). The frame color indicates the directionality: Me3*HIST1*^low^ (blue) and Me3*HIST1*^high^ (red). (**E**) Gene set enrichment analysis of Me3*HIST1*^low^ and Me3*HIST1*^high^ AMLs against CD36_signarure. (**F**) Venn diagram and IGV screen shot showing the unique common gene between the CD36 signature, the NPM1^mut^ DN genes (ALCALAY_DN) and the Me3*HIST1*^low^ UP genes.

Overall, these transcriptomic analyses confirmed the difference in transcriptomes of Me3*HIST1*^low^ and Me3*HIST1*^high^ blasts and highlighted a difference in cell maturation and metabolism that may account for the difference in patient survival between the two groups of patients.

### Me3*HIST1*^high^ phenotype is related to epigenetic signature of NPM1^mut^

Enhancer activity has proven to efficiently demark subgroups of AML patients ^35^. To explore enhancer activity across the two groups, we first defined active (H3K27ac-enriched) and poised (H3K27me3-enriched) enhancers as previously described ^36^ and then analyzed the dynamic of their activity between the two groups of *NPM1*^mut^ AMLs. The main difference in enhancer activity we observed between the two conditions was a gain in H3K27ac (de novo active) in Me3*HIST1*^high^ condition (**Figure 5A**). To assess the H3K27ac influence on DEGs, we crossed enhancers differentially enriched in H3K27ac with the DEGs. Of the 1,028 genes differentially expressed between the two conditions, 103 genes (10%) presented a significant gain or loss of H3K27ac in enhancer region in correlation with the direction of transcriptional deregulation (**Table S4**; **Figure 5B)**. Some of these genes showed also a difference in their TSS region (**Figure S5A)**. The NPM1 mutant protein has been recently shown to amplify leukemic transcriptional programs by maintaining active chromatin ^8^. We investigated whether we could enrich the Me3*HIST1*^high^ signature with *NPM1*^mut^ genes when taking into account the chromatin information. We first crossed our gene lists with previous *NPM1* signature genes (**Table S5**). Consistent with our GSEA, 24 Me3*HIST1*^high^ UP genes (3.6%) were found in either Alacalay_UP or Verhaak_UP (**Figure 5C, left panel)**, whereas only 6 Me3*HIST1*^high^ DN genes were common with Alacalay_UP and/or Verhaak_UP (**Figure S5B**). Interestingly, when considering Me3*HIST1*^high^ UP with a positive correlation of H3K27ac levels in their regulatory regions (Cross RNA/K27ac-expression UP) we enriched the Me3*HIST1*^high^ UP with *NPM1*^mut^ specific genes (up to 11.6%; **Figure 5C**). To precise a direct effect of NPM1^*mut*^ on these genes, we crossed our gene lists with *NPM1*^mut^ chromatin targets identified by ChIP-seq (**Table S4**, REF). We highlighted 4 genes with 2 of them showing correlated change in H3K27ac (**Table S5; Figure 5D)**. IGV visualization confirmed the mRNA and H3K27ac increase at these *NPM1*^mut^ AML genes in Me3*HIST1*^high^ patients (**Figure 5E**).

**Figure 5:**
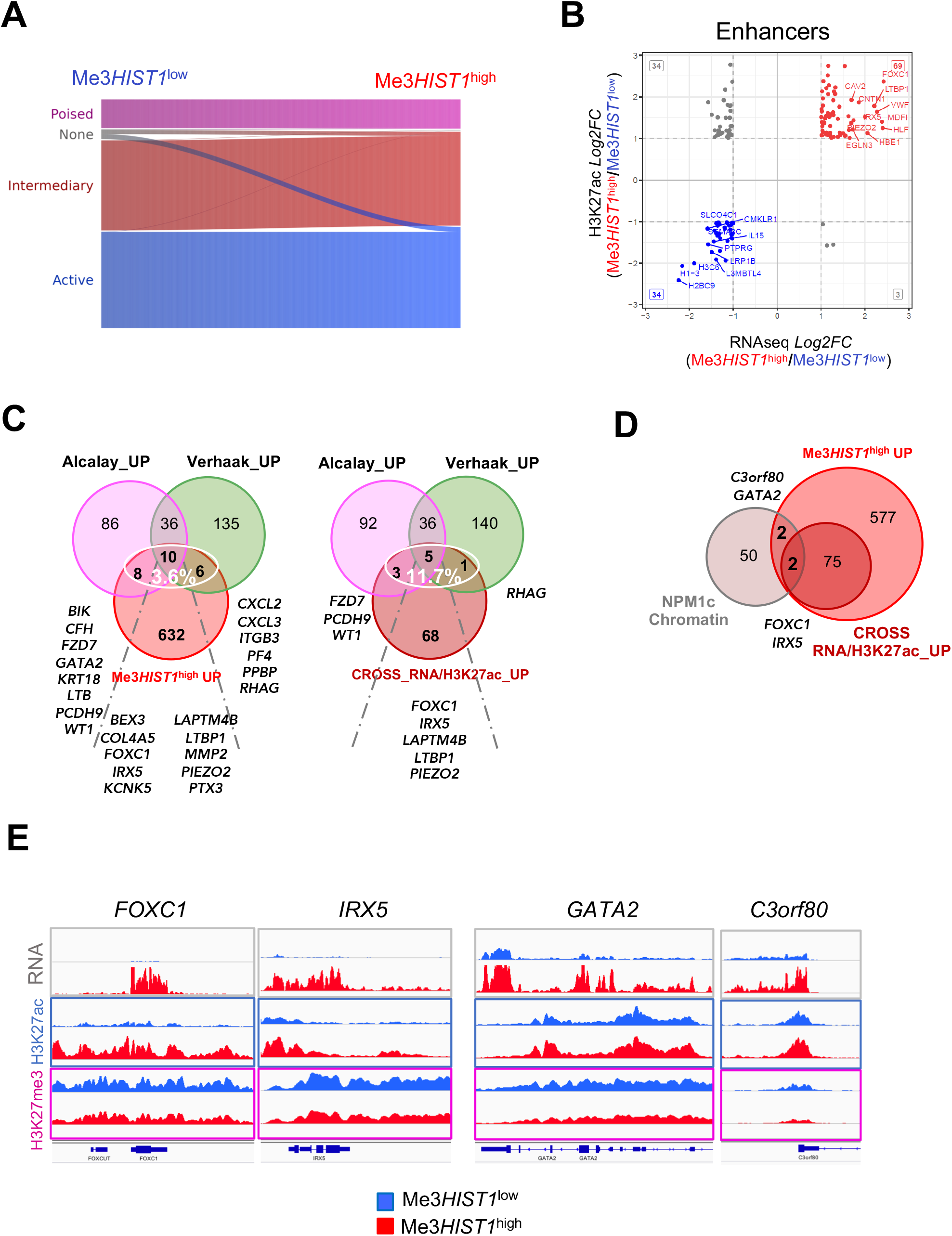
Correlated epigenetic and gene expression between Me3*HIST1*^low^ and Me3*HIST1*^high^ patients. (**A**) Alluvial plot of enhancer activity dynamics between Me3*HIST1*^low^ (left) and Me3*HIST1*^high^ (right) patients. (**B)** Scatter plot of differential H3K27ac level at enhancers (y-axis) and differential gene expression (x-axis) changes between the two subgroups. The top 10 genes in quadrant of similar Log2FC direction are labeled. (**C**) Non-proportional Venn plot to compare genes in NPM1^mut^ UP lists (Alcalay_UP, Verhaak_UP) with Me3*HIST1*^high^ UP genes (left panel) and with _UP and Cross_RNA/H3K27ac_UP gene (right panel). Common genes are listed. Percentage represent the enrichment of Alcalay_UP and Verhaak_UP signatures in our lists of genes. (**E**) Venn plot of NPM1c_chromatin_Targets, Me3*HIST1*^high^ UP and Cross Me3*HIST1*^high^ /H3K27ac_UP gene lists. Common genes are listed. (**F**) mRNA, H3K27ac and H3K27me3 signals at the 4 common loci (NPM1c chromatin-Me3HIST1^high^ UP) in Me3*HIST1*^low^ and Me3*HIST1*^high^ AMLs.

All together, these results suggest that Me3*HIST1*^high^ patients express a specific *NPM1*^mut^ gene signature in relation to a gain of H3K27ac at TSS or Enhancers of these genes and that these enrichments may be due to neomorphic NPM1^mut^ chromatin functions. Our results confirm the interest of using epigenetic information to stratify patients in an already well characterized but heterogeneous AML subgroup and to pinpoint potential driver genes.

## DISCUSSION

Here, we have characterized two groups of CN-*NPM1*^mut^ AML defined by their level of H3K27me3 in part of the *HIST1* locus affecting 11 histone genes. We confirmed the specificity of this H3K27me3 chromatin domain in a fraction of the CN-*NPM1*^mut^ AML patients. Although we still do not know the molecular basis of this difference across the patients, we think that its appearance is tightly linked with the *NPM1* mutation. Indeed, it is predominantly found in *NPM1*mut AML ^16^. NPM1 was recently shown to maintain H3K27me3 chromatin domains throughout DNA replication and to participate in restoring full repressed chromatin states throughout S phase ^37^, its mutation may thus perturbate this function and favor abnormal H3K27me3 deposition. One of the main questions is whether this domain may reflect a differentiation stage that would be caught during the transformation process. Although the two groups are different in term of differentiation genes, with Me3*HIST1*^high^ being more immature and megakaryocytes primed and the Me3*HIST1*^low^ group being more committed towards myeloid differentiation, the absence of H3K27me3 domain differences at the *HIST1* locus through the hematopoietic cells tested would suggest that it is not the case. However, the fact that we detected the same H3K27me3 domain in the MSC and its differentiated lineages (osteoblasts, chondrocytes and adipocytes), which constitute the important components of the hematopoietic microenvironmental niche ^38^, raises the issue of the influence of MSC on leukemias. This influence could take the form of an exchange between leukemia cells an exchange between leukemic cell and MSC through exosomes ^39,40^, although epigenetic information exchange has not been described.

AML has somehow the most robust genetic risk stratification system in the cancer field ^41^. However, due to the heterogeneity and the numerous undetected genetic alterations that have led to some unsatisfactory results, different research groups are developing intracellular signaling or chromatin-based AML classifications that reflect different cell phenotypes and treatment responses ^42,43^. Here, we clearly describe that a large domain of H3K27me3 separates *NPM1*^mut^ AML entity into two groups with quite distinct molecular characteristics, suggesting the stable and reliable nature of this epigenetic biomarker for stratifying AMLs. The two groups have different transcriptomes, epigenetic profiles, and cell surface marker profiles, allowing us to identify two main features that separate these groups. One notable feature is the difference related to metabolism between the two groups of patients; Me3*HIST1*^low^ group being characterized with high CD36 surface expression and reflected with a high expression of one of its effector *KUNY* and higher levels of H3K4me3 in the promoter of enzymatic genes. CD36 is involved in metabolism and signals resistance to chemotherapies ^44,45^, while H3K4me3 regulates promoter-proximal pause release and facilitates gene expression ^8^. These differences in surface molecules, reflected in enzymatic activities and metabolism, may provide a molecular explanation for the phenotype ^16^ and re-enforce the link between epigenetics and metabolism in the physiopathology of AML.

The second notable feature is the more pronounced *NPM1*^mut^ phenotype in the Me3*HIST1*^high^ leukemia. Since low CD34 expression is a characteristic of NPM1^mut^ blasts, this translates into even lower CD34 expression on the surface of Me3*HIST1*^high^ blasts compared to Me3*HIST1*^low^ blasts. It also translates into the expression of *NPM1*^mut^ signature that is enriched in Me3*HIST1*^high^ AML. From a molecular point of view, this more pronounced expression could have epigenetic origins. Part of the *HIST1* locus that is enriched with the H3K27me3 delineates a TAD domain ^46^, which could influence the main epigenetic changes observed between the two groups, involving H3K27ac at enhancer regions. 11% of the genes presenting H3K27ac/gene expression differences between the two conditions have recently described as being regulated by *NPM1*^mut 9^. This suggests that carrying the Me3*HIST1*^high^ marker may drive stronger expression of NPM1^mut^-associated genes via enhancer activation. Although we do not know the mechanism underlying this increase in enhancer activity in Me3*HIST1*^high^ patients, it is interesting to note that the two genes FOXC1 et IRX5, which check all the boxes (direct targets of *NPM1*^mut^, strong correlation of H3K27ac levels and expression in Me3*HIST1*^high^) are coregulated with GATA2 in AMLs. This coregulation may pass through the TAD domain associated with an intron of the *FTO* (fat mass and obesity-associated) gene on chromosome 16^47^. One of the interesting candidates is the high expression, coupled with H3K27ac accumulation, in the transcription factor *FOXC1* in Me3*HIST1*^high^. FOXC1 is a critical regulator of HSC niche development and maintenance ^48^ and is usually associated with poor prognosis. However, in certain cancer types, elevated *FOXC1* expression, has been shown to be a predictor of favorable prognosis ^49^. Activity of FOXC1 may be linked to *HIST1* locus and in the context of *NPM1*^mut^ would predict favorable prognosis by improving the tumor microenvironment.

In conclusion, our work demonstrates the importance of using epigenetic information to stratify patients in an already well-characterized but heterogeneous AML subgroup. Our study also shows how an epigenetic regulation can influence physiological differences between AML blasts and help predict response to treatment.

## Acknowledgements

We would like to thank all the patients who donated samples for this study. The authors wish to thank Jihane Pakradouni for helping selecting patient samples, the CIBI and DISC platforms for computational analyses and support, the Helixio platform for the RNA sequencing and the IPC genomic sequencing facility for the NGS for ChIP-seq.

